# The histone demethylase KDM5 is essential for larval growth in *Drosophila*

**DOI:** 10.1101/297804

**Authors:** Coralie Drelon, Helen M. Belalcazar, Julie Secombe

## Abstract

Regulated gene expression is necessary for developmental and homeostatic processes. The KDM5 family of proteins are histone H3 lysine 4 demethylases that can regulate transcription through both demethylase-dependent and independent mechanisms. While loss and overexpression of KDM5 proteins are linked to intellectual disability and cancer, respectively, their normal developmental functions remain less characterized. *Drosophila melanogaster* provides an ideal system to investigate KDM5 function, as it encodes a single ortholog in contrast to the four paralogs found in mammalian cells. To examine the consequences of complete loss of KDM5, we generated a null allele of *Drosophila kdm5*, also known as *little imaginal discs* (*lid*), and show that it is essential for development. Animals lacking KDM5 die during late pupal development but show a dramatically delayed larval development that coincides with decreased proliferation and increased cell death in imaginal discs. Interestingly, this developmental delay is independent of the well-characterized Jumonji C (JmjC) domain-encoded histone demethylase activity and plant homedomain (PHD) motif-mediated chromatin binding activities of KDM5, suggesting key functions for less characterized domains. Consistent with the phenotypes observed, transcriptome analyses of *kdm5* null mutant wing imaginal discs revealed the dysregulation of genes involved in several cellular processes, including cell cycle progression and DNA repair. Together, our data provide the first description of complete loss of KDM5 function in a metazoan and offer an invaluable tool for defining the biological activities of KDM5 family proteins.

## Introduction

Regulated gene expression is essential for growth and cell fate decisions critical to development, in addition to the maintenance of tissues and organs during adulthood. Changes to chromatin, the structure that includes DNA and its associated histone proteins, is one key mechanism by which transcription is regulated (Swygert and Peterson 2014). Histones are extensively decorated by covalent modifications that can impact chromatin compaction to influence transcription factor binding and/or affect the recruitment of proteins that recognize specific histone modifications to activate or repress promoter activity (Rothbart and Strahl 2014). One family of transcriptional regulators that both recognizes and enzymatically modifies chromatin is the lysine demethylase 5 (KDM5) family of evolutionarily conserved histone demethylases. Mammalian cells encode four KDM5 paralogs, KDM5A, KDM5B, KDM5C and KDM5D, whereas organisms with smaller genomes such as *Drosophila* and *C. elegans* have a single KDM5 protein.

The driving force behind defining the physiological functions of KDM5 proteins stems from the observation that dysregulation of *KDM5* family genes is linked to human diseases. Specifically, loss of function mutations in *KDM5A*, *KDM5B* and *KDM5C* are found in patients with intellectual disability (ID), link KDM5 function to neuronal development or function (Vallianatos and Iwase 2015). Altered expression of *KDM5* family genes is also implicated in cancer. KDM5A and KDM5B, both of which are encoded on autosomal chromosomes, are candidate oncoproteins (Blair *et al*. 2011). They are overexpressed in a number of different cancers including breast, ovarian and lung, and their expression correlates with both proliferation rate and propensity for metastatic invasion (Hayami *et al*. 2010; Hou *et al*. 2012; Teng *et al*. 2013; Yamamoto *et al*. 2014; Wang *et al*. 2015; Feng *et al*. 2017; Huang *et al*. 2018). The link between the X-linked *KDM5C* gene and cancer is less straight-forward, as it has both oncogenic and tumor suppressive activities in tissue-dependent manner (Harmeyer *et al*. 2017). The Y-linked *KDM5D* gene is a tumor suppressor and prognostic indicator of poor outcome in cases of prostate cancer (Komura *et al*. 2016; Li *et al*. 2016). Together, these data suggest that KDM5 proteins can play context-dependent roles in neuronal development, cell survival and proliferation.

Although the molecular mechanisms by which dysregulation of KDM5 proteins causes disease remains unknown, it is likely that it is related to their roles in transcriptional regulation. The Jumonji C (JmjC) catalytic domain is the most extensively characterized domain of KDM5 and is responsible for demethylating histone H3 that is trimethylated at lysine 4 (H3K4me3) (Klose and Zhang 2007). High levels of H3K4me3 are found most predominantly surrounding the transcription start site (TSS) regions of actively expressed genes (Santos-Rosa *et al*. 2002). Thus, the demethylase activity of KDM5 is thought to lead to transcriptional repression. The role of this enzymatic activity in disease remains only partially resolved. While missense mutations in KDM5C associated with intellectual disability can impair demethylase activity *in vitro*, this is not universally true (Tahiliani *et al*. 2007; Vallianatos and Iwase 2015). In addition, while the proliferation and survival of some KDM5 overexpressing cancers rely on enzymatic activity (Teng *et al*. 2013), others do not (Cao *et al*. 2014), implicating key activities for other domains. One example of this is the C-terminal PHD domain (PHD3) that binds to H3K4me2/3 (Wang *et al*. 2009; Li *et al*. 2010; Klein *et al*. 2014). This domain is involved in the activation of genes necessary for metabolic and mitochondrial functions (Liu and Secombe 2015). The importance of this PHD motif to regulated gene expression is emphasized by the observation that this domain can cause leukemia when fused to the nuclear pore protein NUP98 (Van zutven *et al*. 2006; Wang *et al*. 2009). Thus, KDM5 proteins can act through distinct mechanisms to affect gene expression, but the molecular links between these transcriptional activities and disease are not well understood.

Despite a growing body of evidence linking loss or gain of KDM5 proteins to human disease, the normal developmental functions of KDM5 proteins remain unclear. To begin addressing the normal functions of KDM5 *in vivo*, the genetic advantages of model organisms such as mice, flies and worms have been employed. Significantly, some phenotypes observed are consistent with the clinical features found in patients with altered levels of KDM5 proteins. For example, *Kdm5C* knockout mice show cognitive phenotypes, consistent with the mutations observed in human intellectual disability patients (Iwase *et al*. 2016; Scandaglia *et al*. 2017). Likewise, hypomorphic mutations in the *C. elegans* KDM5 ortholog *rbr-2* result in axonal guidance defects (Lussi *et al*. 2016; Mariani *et al*. 2016). In addition, while *Kdm5A* knockout mice are viable, cultured MEFS from these animals show reduced proliferation, implicating KDM5A as a positive regulator of cell cycle progression (Lin *et al*. 2011). *Kdm5B* and *Kdm5C* knockout mice also survive but they are smaller than their littermates, suggesting roles for these KDM5 family proteins in growth regulation (Zou *et al*. 2014; Iwase *et al*. 2016). Significantly, mouse studies carried out to-date have been complicated by phenotypes that are dependent on strain background, raising questions regarding whether phenotypes observed are specifically due to loss of the *Kdm5* gene being examined (CATCHPOLE *et al*. 2011; Albert *et al*. 2013). Moreover, there is evidence that loss of one *KDM5* family gene can cause compensatory upregulation of another family member, potentially masking additional processes that rely on KDM5 function (Jensen *et al*. 2010; Zou *et al*. 2014).

*Drosophila* encodes a single *kdm5* gene, also known as *little imaginal discs* (*lid*). Reducing the levels of KDM5 through hypomorphic alleles or by ubiquitous RNAi-mediated knockdown causes semi-lethality (Gildea *et al*. 2000; Li *et al*. 2010; Lloret-Llinares *et al*. 2012) suggesting that in the absence of paralogs, *kdm5* may be necessary for development. However, analyses of the phenotypes caused by a complete absence of KDM5 have been hampered by the absence of a *kdm5* null allele. Here we describe the generation of the amorphic allele *kdm5*^*140*^ and show that KDM5 is essential for development. Specifically, we demonstrate that loss of KDM5 causes a dramatic developmental delay and pupal lethality that correlates with decreased rates of proliferation and increased cell death in larval imaginal discs. Transcriptome analyses of *kdm5*^*140*^ wing imaginal discs showed that loss of KDM5 alters the expression of genes involved in a number of cellular processes, including cell cycle progression and DNA dam@age repair. Consistent with this, *kdm5*^*140*^ mutants are sensitive to DNA damaging agents. Together, our results demonstrate that KDM5 is a key regulator of larval growth and provides an invaluable tool to further dissect the biological roles of KDM5 proteins.

## Materials and methods

### Fly strains and husbandry

All flies were kept at 25°C on standard food with 60% humidity and a 12 hour light/dark cycle. The *kdm5:HA*^*WT*^ and *kdm5:HA*^*JmjC\**^ transgenes are published (NAVARRO-COSTA *et al*. 2016; Zamurrad *et al*. 2018). Briefly, they are 11kb constructs that include the entire *kdm5* genomic region with the addition of three HA tags at the 3’ end of the open reading frame. Transgenes were inserted into the attP site at 86Fb and the *white* and *RFP* cassette removed using germline expression of the Cre recombinase (Bischof *et al*. 2007). The *kdm5:HA*^*W1771A*^ transgene is identical to *kdm5:HA*^*WT*^ except for a codon change that alters tryptophan 1771 to alanine. The *kdm5:HA*^*W1771A*^ transgene and corresponding control wildtype transgene were inserted into the attP site at 68A4 on chromosome III (generated at the “Rainbow transgenic flies”)(Groth *et al*. 2004). This *kdm5:HA*^*W1771A*^ construct is similar to a previously published transgene that was not HA tagged, did not contain *kdm5* introns and was randomly integrated into the *Drosophila* genome (Liu and Secombe 2015). The homozygous viable *kdm5*^*NP4707*^ strain was obtained from the Kyoto Stock Center (Kyoto Institute of Technology; stock #104754) and *kdm5*^*140*^ was generated by imprecise excision of this *P* element using delta 2-3 transposase. The breakpoints of *kdm5*^*140*^ were molecularly mapped by genomic PCR and subsequent sequencing (Figure S1). All other strains were obtained from the Bloomington *Drosophila* stock center.

### Developmental timing analyses

10 females and males were crossed and allowed to lay eggs for 24 hours in food vials, with parental flies being subsequently removed. The number of animals that had pupariated was scored twice per day. Experiments were carried out in biological triplicate.

### γ-irradiation

Third-instar larvae were exposed to a 40 Gy dose of γ-radiation using a 137Cs irradiator (Shepherd Mark I Irradiator). After 4 hours at 25°C, wings discs were dissected and used for Dcp-1 immunostaining. Fluorescence intensity of Dcp1 staining was quantified using ImageJ Software and divided by the area of that wing disc to provide a ratio of intensity/area.

### Larval food ingestion analyses

To monitor food intake, adult flies were allowed to lay eggs on fly food containing 0.05% bromophenol blue. Larvae were removed from the food 36 hours later and examined by light microscopy.

### Immunostaining

For immunostaining, wing imaginal discs from 3^rd^ instar larvae were dissected in 1X PBS and fixed in 4% paraformaldehyde for 30 minutes at room temperature. They were then washed 3 times with PBST (1X PBS, 0.2% Triton), blocked 1 hour at 4°C in 0.1% BSA, followed by the incubation with the primary antibody overnight at 4°C. Anti-pH3 (Cell signaling #9701) and anti-Dcp1 (Cell signaling #9578) were used at 1/50 and 1/100, respectively. For pupal wing staining, white prepupae were picked and aged until 28-30hrs APF (after puparium formation) at 25°C before fixing in 4% paraformaldehyde overnight at 4°C. Samples were then washed 3 times with PBST, pupal wings dissected, blocked 1 hour at 4°C with 0.1% BSA/PBST and incubated with the anti-22C10 (DSHB, University of Iowa, 1/50) overnight at 4°C. Secondary antibodies were obtained from Invitrogen. Wings were mounted in Vectashield and imaged on a Zeiss Axio Imager 2 microscope. For quantification, the number of pH3 and Dcp1 positive cells in the wing pouch were counted using Image J software.

### Western blot

Western blots were carried out as previously described using LiCOR (Liu and Secombe 2015). Antibodies used were anti-pH3 (Cell signaling #9701, 1/1000), anti-histone H3 (Active Motif #39763 or #39163, 1/5000), anti-alpha Tubulin (Developmental Studies Hybridoma Bank, University of Iowa; 1:5000). The rabbit polyclonal KDM5 antibody was raised to amino acids 1418 to 1760 and has been previously published (Secombe *et al*. 2007).

### Translation quantification

Translation levels in wing imaginal discs was quantified as previously described (Deliu *et al*. 2017). Briefly, wing discs from 3^rd^ instar larvae were dissected and incubated in 5 micrograms/ml puromycin (Sigma) or puromycin plus 10 micrograms/ml cycloheximide (Sigma). Puromycin levels in wing discs were then assayed by Western blot with anti-puromycin (3RH11; 1:1000; Kerafast). Anti-histone H3 (Active Motif #39763; 1:1000) was used as a loading control. Quantification was carried out using LiCOR software by determining the intensity of all puromycin labeled proteins between 14 and 200kDa and dividing this by the intensity of the histone H3 load control.

### RNA-seq

RNA-seq was carried out at the New York Genome Center. RNA was prepared in biological triplicate from wildtype (*kdm5*^*140*^; *kdm5:HA*^*WT*^; referred to as *kdm5*^*WT*^), and *kdm5*^*140*^ 3^rd^ instar mutant larval wing discs matched for developmental age using Trizol and RNAeasy (Qiagen). An equal mix of male and female larvae were used. RNA sequencing libraries were prepared using the TruSeq Stranded mRNA Library Preparation Kit in accordance with the manufacturer’s instructions. Briefly, 500ng of total RNA was used for purification and fragmentation of mRNA. Purified mRNA underwent first and second strand cDNA synthesis. cDNA was then adenylated, ligated to Illumina sequencing adapters, and amplified by PCR (8 cycles). Final libraries were evaluated using fluorescent-based assays including PicoGreen (Life Technologies) or Qubit Fluorometer (Invitrogen) and Fragment Analyzer (Advanced Analytics) or BioAnalyzer (Agilent 2100) and were sequenced on an Illumina HiSeq2500 sequencer (v4 chemistry) using 2 × 50bp cycles. Alignment of raw reads was carried out using STAR aligner, normalized and differential expression determined with DESeq2. The accession number for the RNA-seq data described here is GSE109201. Volcano plot showing dysregulated genes was generated with ggplot2 package in R.

### Real-time PCR

Real-time PCR was carried out as previously described (Liu and Secombe 2015) using cDNA from 3^rd^ instar larval wing imaginal discs. Primer sequences are provided in Table S1.

### Statistical analyses

All experiments were done in biological triplicate (minimum) and Ns are provided for each experiment. Fisher’s exact test was carried out the program R (v3.3.2). Student’s t-test, chi-squared and Wilcoxen rank-sum tests were carried out using GraphPad Prism version 7.00 (GraphPad Software, La Jolla CA).

### Data availability

A list of differential expressed genes (and log2 fold change) observed in *kdm5*^*140*^ compared to wildtype is provided in Table S2 (5% FDR). Table S2 also includes data from KDM5 ChIP-seq that is published and publically available from adult flies (GSE70591) (Liu and Secombe 2015) and wing imaginal discs (GSE27081)(Lloret-Llinares *et al*. 2012). Table S3 shows genes dysregulated in *kdm5*^*140*^ RNA-seq and *kdm5*^*10424*^ wing disc microarray data (GSZ53881) (Liu *et al*. 2014). Table S4 shows genes dysregulated in and *kdm5*^*140*^ RNA-seq and S2 cell KDM5 knockdown RNA-seq (GSE68775) (Gajan *et al*. 2016). The overlap between genes affected in *kdm5*^*140*^ RNA-seq, *kdm5*^*10424*^ microarray, S2 cell knockdown RNA-seq and *kdm5* knockdown wing disc microarray (GSE27081) (Lloret-Llinares *et al*. 2012) is provided in Figure S2.

## Results

### *kdm5* is an essential gene required for developmental timing

Existing hypomorphic *P* element alleles of *kdm5* such as *kdm5*^*10424*^ and *kdm5*^*K06801*^ are 95% lethal (Gildea *et al*. 2000; Secombe *et al*. 2007) (Figure 1A, 1B). *kdm5* mutant flies that do eclose are morphologically normal, but are short-lived (Liu and Secombe 2015). Because the effects of complete loss of KDM5 remain unknown, we generated a null allele named *kdm5*^*140*^ by imprecise excision of a homozygous viable *P* element insertion in the *kdm5* promoter region (*kdm5*^*NP4707*^; Figure 1B). Molecular mapping demonstrated that this allele deletes three of the four coding exons of the *kdm5* gene, including the start codon (Figure 1B; Figure S1). Real-time PCR analyses of *kdm5*^*140*^ homozygous mutant larvae demonstrated a complete loss of full length *kdm5* transcript compared to the genetically similar wildtype strain *w*^*1118*^, although a partial 3’ end transcript remains (Figure 1C, D). Despite this truncated transcript having an open reading frame with the potential to encode a 51kDa protein, Western blot demonstrated that no full length or truncated protein(s) were present in *kdm5*^*140*^ animals (Figure 1E). *kdm5*^*140*^ null mutants are 100% lethal (Figure 1A). To demonstrate that this lethality was due specifically to the loss of KDM5, we utilized a previously generated genomic rescue transgene containing a HA-tagged form of the *kdm5* locus (Figure 1F)(Navarro-Costa *et al*. 2016; Zamurrad *et al*. 2018). Because the expression of this transgene is driven by the endogenous *kdm5* promoter region, it is expressed at wildtype levels and is able to rescue the lethality of the *kdm5*^*140*^ allele (Figure 1G, H). *kdm5* is therefore an essential gene in *Drosophila*.

**Figure 1:**
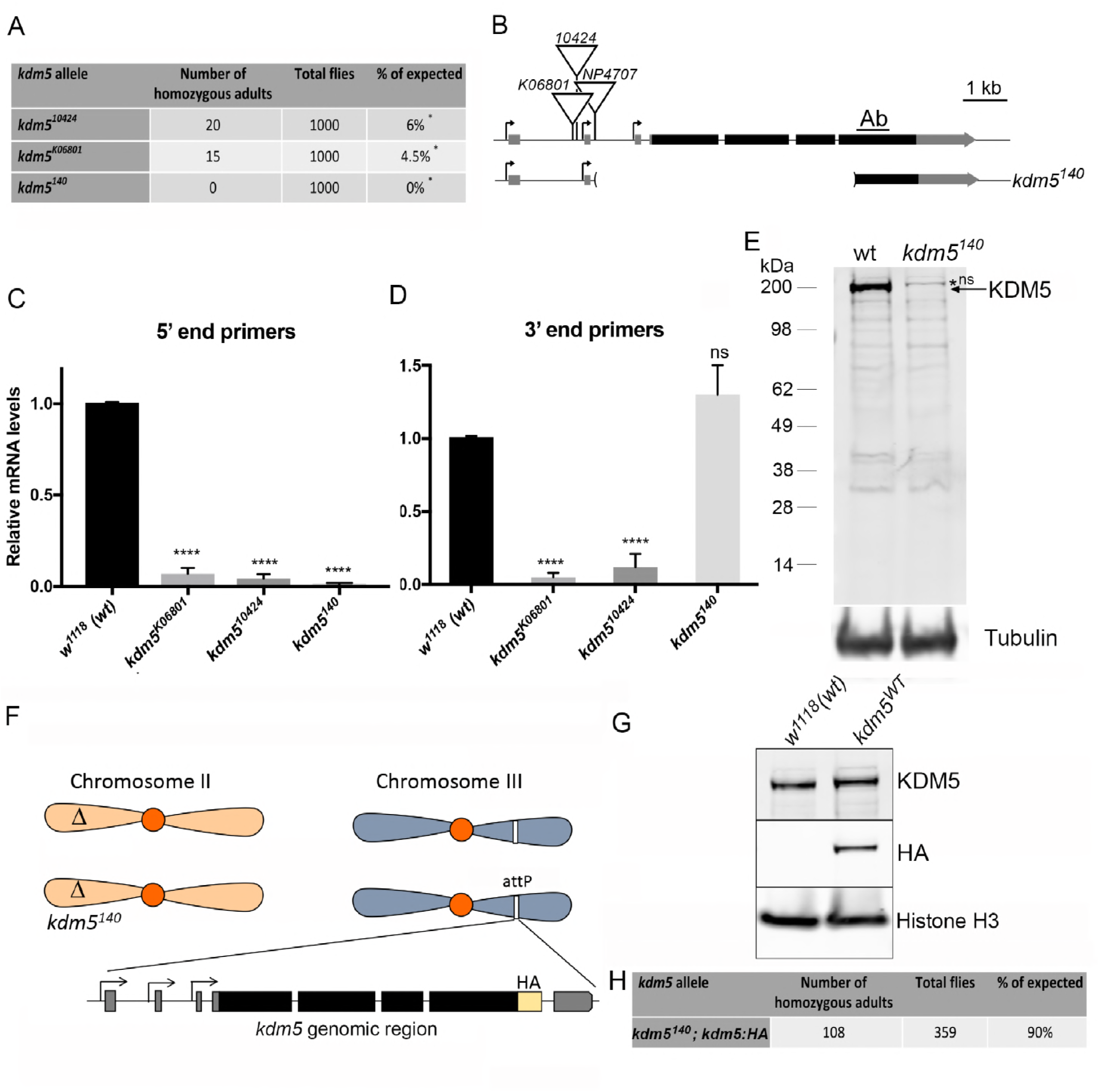
A *kdm5*^*140*^ null allele demonstrates that KDM5 is essential in *Drosophila*. (A) Lethality of *kdm5*^*K06801*^, *kdm5*^*10424*^ and *kdm5*^*140*^ homozygous mutant animals generated from a cross between five female and five male heterozygous parents balanced using CyO-GFP. The column labeled total flies indicates the number of progeny (adult) flies scored from at least three independent crosses. Expected number of progeny is based on Mendelian frequencies and taking into account the lethality of CyO homozygotes, i.e 33% of total adult flies. * *p*<0.01 (chi-squared test). (B) Position of the *NP4707*, *10424* and *K06801 P* element insertions and molecular mapping of the *kdm5*^*140*^ deletion. Ab indicates the region used to generated the rabbit polyclonal anti-KDM5 antibody (Secombe *et al*. 2007). (C) RT-PCR using primers to the 5’ end of the gene using RNA from whole 3^rd^ instar larvae. Animals homozygous for *kdm5*^*K06801*^ or *kdm5*^*10424*^ show low levels of transcript while *kdm5*^*140*^ shows none. *kdm5* mRNA normalized to wildtype (*w*^*1118*^) using *rp49*. **** *p*<0.0001. (D) RT-PCR using primers to the 3’ end of the gene using RNA from whole 3^rd^ instar larvae. *kdm5*^*140*^ has wildtype levels of the 3’ end of the transcript. **** *p*<0.0001. ns = not significant. (E) Western from wildtype (*w*^*1118*^) and *kdm5*^*140*^ homozygous mutant wing imaginal discs showing KDM5 and alpha tubulin. *kdm5*^*140*^ animals have no detectable full length or truncated KDM5. *ns indicates non-specific band. (F) Schematic of strain genotype for rescue of *kdm5*^*140*^ with a genomic rescue transgene. Flies are homozygous for the *kdm5*^*140*^ mutation on the 2^nd^ chromosome and homozygous for an 11kb genomic rescue transgene on the 3^rd^ chromosome. (G) Western blot showing KDM5 protein levels from 3^rd^ instar larval wing imaginal discs from wildtype (*w*^*1118*^) and *kdm5*^*140*^ homozygotes that also have two copies of the *kdm5:HA* genomic rescue transgene. Anti-KDM5 (top), anti-HA (middle) and anti-histone H3 loading control (bottom). (H) *kdm5*^*140*^ lethality is rescued by a transgene encoding the *kdm5* locus. These data were generated by crossing female and male flies heterozygous for *kdm5*^*140*^ and homozygous the wildtype genomic rescue transgene (intercross of *kdm5*^*140*^/CyO-GFP; *kdm5:HA*/*kdm5:HA* males and females).

To understand the basis for KDM5’s essential functions, we quantified the developmental timing of *kdm5*^*140*^ homozygous mutants. To ensure that we were examining defects due to the loss of KDM5, these and all subsequent experiments utilized a wildtype control strain in which the *kdm5*^*140*^ null mutation was rescued by two copies of the *kdm5* genomic rescue transgene (*kdm5*^*WT*^). Whereas *w*^*1118*^, *kdm5*^*WT*^ and *kdm5*^*140*^ heterozygous animals have indistinguishable developmental timings and took an average of 6.8 days for 50% of animals to pupariate, *kdm5*^*140*^ homozygous mutant larvae took an average of 12 days (Figure 2A). To rule out the possibility that *kdm5*^*140*^ mutants were developmentally delayed due to decreased food consumption, we fed first instar larvae food containing bromophenol blue. As shown in Figure 2B, *kdm5*^*WT*^ and *kdm5*^*140*^ mutant larvae show similar levels of ingested dye, demonstrating that KDM5 is not required for feeding. *kdm5*^*140*^ mutant animals took longer to progress through larval development, weighing less and having significantly smaller imaginal discs compared to age-matched control larvae (Figure 2C-F). Despite this delay, final larval, imaginal disc and pupal size of *kdm5*^*140*^ mutants were normal (Figure 2G-J). Indeed, *kdm5*^*140*^ mutant pharate adults appear to have morphologically normal heads, thoraces, legs and abdominal segmentation, but fail to eclose (Figure 2K-M).

**Figure 2:**
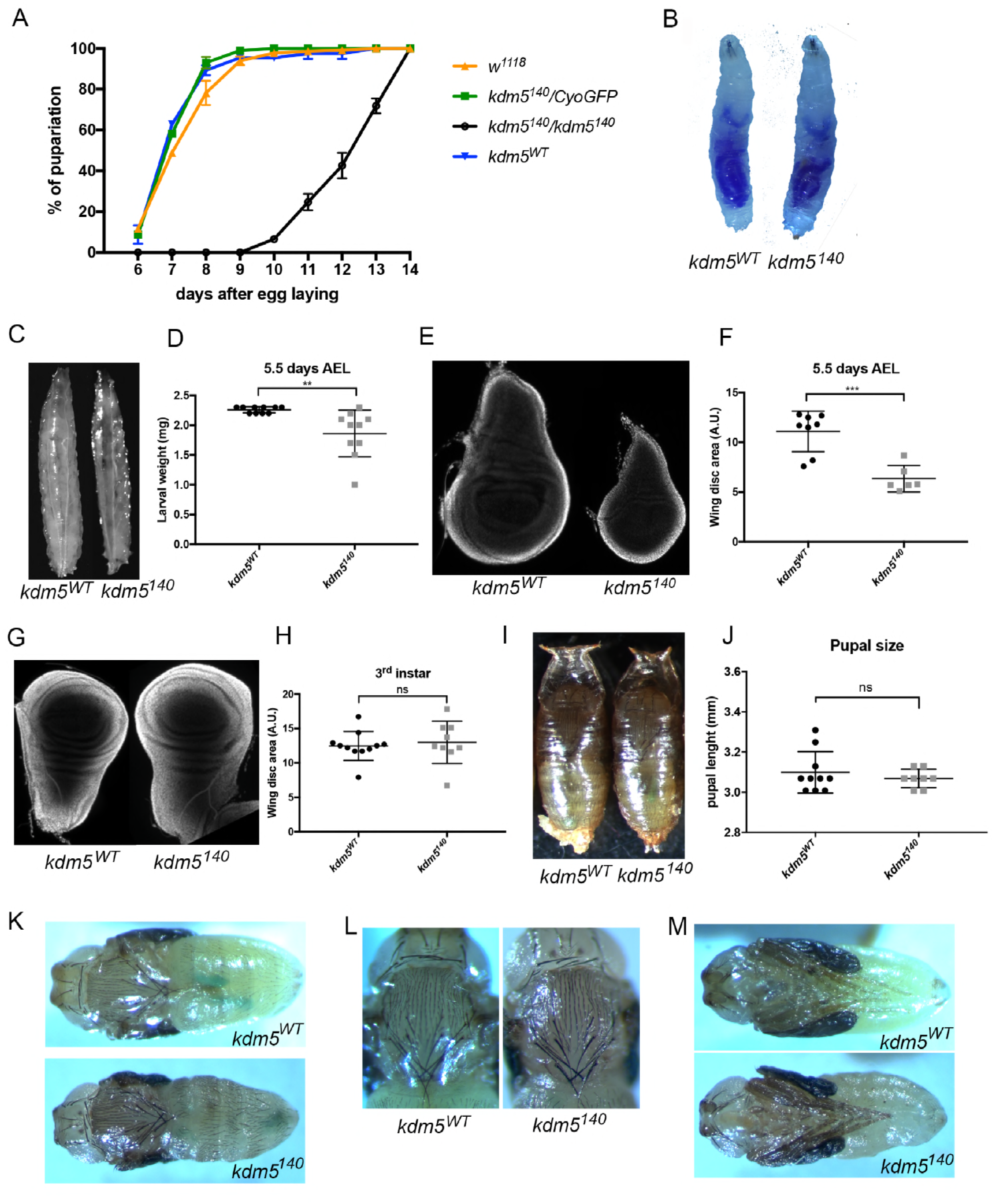
*kdm5*^*140*^ null mutants show a developmental delay and pupal lethality. (A) Time taken for animals to pupariate for *w*^*1118*^ (N=221), *kdm5*^*WT*^ (N=82), *kdm5*^*140*^ heterozygous (*kdm5*^*140*^/CyO-GFP; N=184), or *kdm5*^*140*^ homozygous mutant (N=49). (B) *kdm5*^*WT*^ and *kdm5*^*140*^ mutant larvae fed food containing the dye bromophenol blue. (C) *kdm5*^*WT*^ and *kdm5*^*140*^ mutant larvae at 5.5 days after egg laying (AEL). (D) Weight in milligrams of *kdm5*^*WT*^ (N=10) and *kdm5*^*140*^ mutant (N=10) larvae at 5.5 days. ** *p*=0.005 (E) Wing imaginal discs from *kdm5*^*WT*^ or *kdm5*^*140*^ mutants at 5.5 days AEL. (F) Quantification of the size of *kdm5*^*WT*^ (N=8) and *kdm5*^*140*^ mutant (N=6) wing discs at 5.5 days AEL. *** *p*=0.0003 (G) Wing imaginal discs from *kdm5*^*WT*^ or *kdm5*^*140*^ mutants at wandering 3^rd^ instar larval stage (5.5 days for *kdm5*^*WT*^, 10 days for *kdm5*^*140*^). (H) Quantification of wing disc size (area) of wing imaginal discs from *kdm5*^*WT*^ (N=11) or *kdm5*^*140*^ mutants (N=9) at wandering 3^rd^ instar larval stage. ns = not significant. (I) 9-day-old *kdm5*^*WT*^ and 14-day-old *kdm5*^*140*^ mutant pupae. (J) Quantification of pupal final size for *kdm5*^*WT*^ (N=10) and *kdm5*^*140*^ mutants (N=7). ns = not significant. (K) Dorsal view of 9-day-old *kdm5*^*WT*^ (top) and 14-day-old *kdm5*^*140*^ mutant (bottom) pupae dissected from their pupal case. (L) Thorax and head of *kdm5*^*WT*^ (left) and *kdm5*^*140*^ mutant (right) pupae. (M) Ventral view of *kdm5*^*WT*^ (top) and *kdm5*^*140*^ mutant (bottom) pupae dissected from their pupal case showing normal morphology.

Two of the most characterized functions of KDM5 are its JmjC domain-encoded demethylase activity that removes H3K4me3, and its C-terminal PHD motif that binds to H3K4me2/3 (PHD3) (Eissenberg *et al*. 2007; Lee *et al*. 2007; Secombe and Eisenman 2007; Liu *et al*. 2014). While both domains involve the same chromatin modification, H3K4me3, they likely regulate transcription independently of each other (Liu and Secombe 2015). We therefore tested whether the developmental delay phenotype observed in *kdm5*^*140*^ animals was dependent on the activity of either of these domains. To do this, we utilized transgenes encoding point mutation forms of KDM5 that abolish the function of JmjC or PHD3 domains that are expressed under endogenous control (*kdm5*^*JmjC\**^ and *kdm5*^*W1771A*^; Figure 3A) (Liu and Secombe 2015; Navarro-Costa *et al*. 2016). By crossing these transgenes into the *kdm5*^*140*^ null allele background, the mutant form is the sole source of KDM5, which are expressed at wildtype levels (Figure 3B). Compared to *kdm5*^*WT*^, *kdm5*^*JmjC\**^ and *kdm5*^*W1771A*^ mutant animals took an average of 6.75 and 8.25 days, respectively, for 50% of animals to pupariate compared to 6.6 days for wildtype (Figure 3C). The developmental time for the *kdm5*^*JmjC\**^ strain was indistinguishable from *kdm5*^*WT*^, demonstrating that the developmental delay observed in *kdm5*^*140*^ mutant animals is independent of its demethylase activity. Although *kdm5*^*W1771A*^ flies took a day longer to pupariate than *kdm5*^*WT*^ animals (*p*=0.002), this was still significantly faster than the delay of five days observed in *kdm5*^*140*^ mutant animals (*p*<0.0001). Thus, while chromatin binding may contribute to the developmental growth functions of KDM5, other activities are also critical.

**Figure 3:**
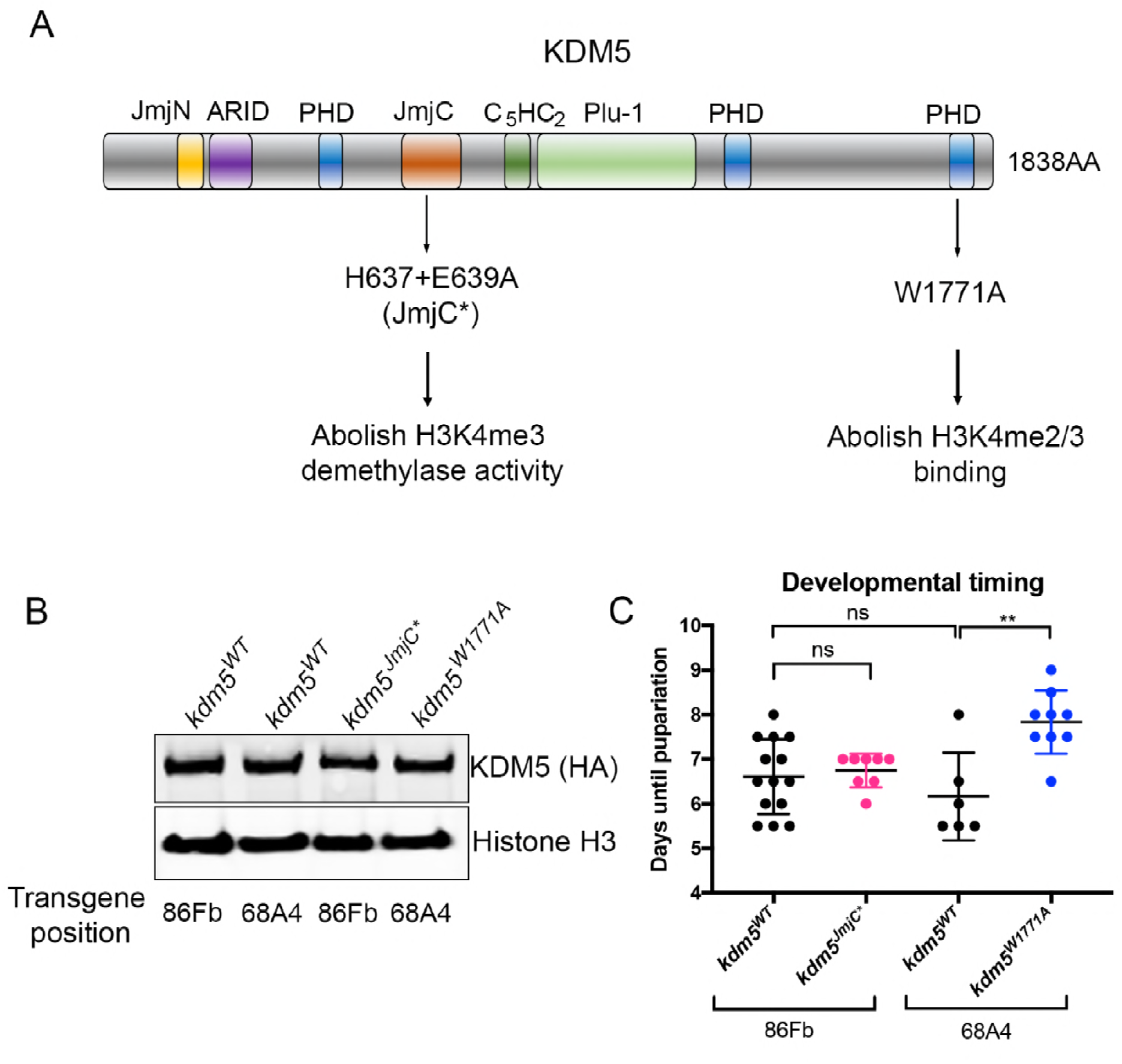
KDM5-mediated developmental delay is independent of its H3K4me3 removal or binding activities. (A) Schematic of the KDM5 protein showing the domain structure and the location of the JmjC* point mutations that abolish demethylase activity and the W1771A point mutation in the C-terminal PHD motif that prevents binding to H3K4me3 (Li *et al*. 2010). The *kdm5*^*JmjC\**^ genomic transgene is inserted at the attP site at 86Fb while the *kdm5*^*W1771A*^ transgene is located at the attP site at 68A4. Each mutant strain therefore has a separate control *kdm5*^*WT*^ strain with a matching insertion of the wildtype *kdm5* genomic region. (B) Western blot showing wildtype expression of KDM5 (using anti-HA; top) in *kdm5*^*JmjC\**^ and *kdm5*^*W1771A*^ wing imaginal discs. The *kdm5*^*WT*^ strain at 86Fb is a control for *kdm5*^*JmjC\**^ while the wildtype insertion at 68A4 is the control for *kdm5*^*W1771A*^. (C) Time for 50% of *kdm5*^*WT*^ (86Fb; N=415), *kdm5*^*JmjC\**^ (86Fb; N=66), *kdm5*^*WT*^ (68A4; N=379) *kdm5*^*W1771A*^ (68A4; N=126) to pupariate (T^1^/_2_). Each data point represents animals counted from an independent cross. N values represent the total number of animals scored. ns = not significant. ** *p*=0.002.

### KDM5 is required for optimal wing disc cell proliferation and survival

The delayed development observed in *kdm5*^*140*^ mutants could, at least in part, be due to slowed proliferation of imaginal disc cells that are the precursors to adult structures. To examine this, we quantified the number of mitotic cells in *kdm5*^*140*^ mutant 3^rd^ instar larval wing imaginal discs using an anti-phospho-histone H3 (pH3) antibody. *kdm5*^*140*^ mutant discs showed ~25% fewer proliferating cells compared to control discs at the same developmental stage (Figure 4A, B). This decreased wing disc proliferation is supported by Western blot analyses demonstrating significantly reduced levels of pH3 (Figure 4C, D). To test whether the reduced proliferation was an indirect consequence of a general cell growth defect, we directly measured levels of translation in *kdm5*^*WT*^ and *kdm5*^*140*^ larvae through incorporation of the tRNA analog puromycin (Deliu *et al*. 2017). Western blot analyses to detect puromycin-labeled proteins showed that wing discs lacking KDM5 have translation rates that were indistinguishable from wildtype (Figure 4E, F).

**Figure 4:**
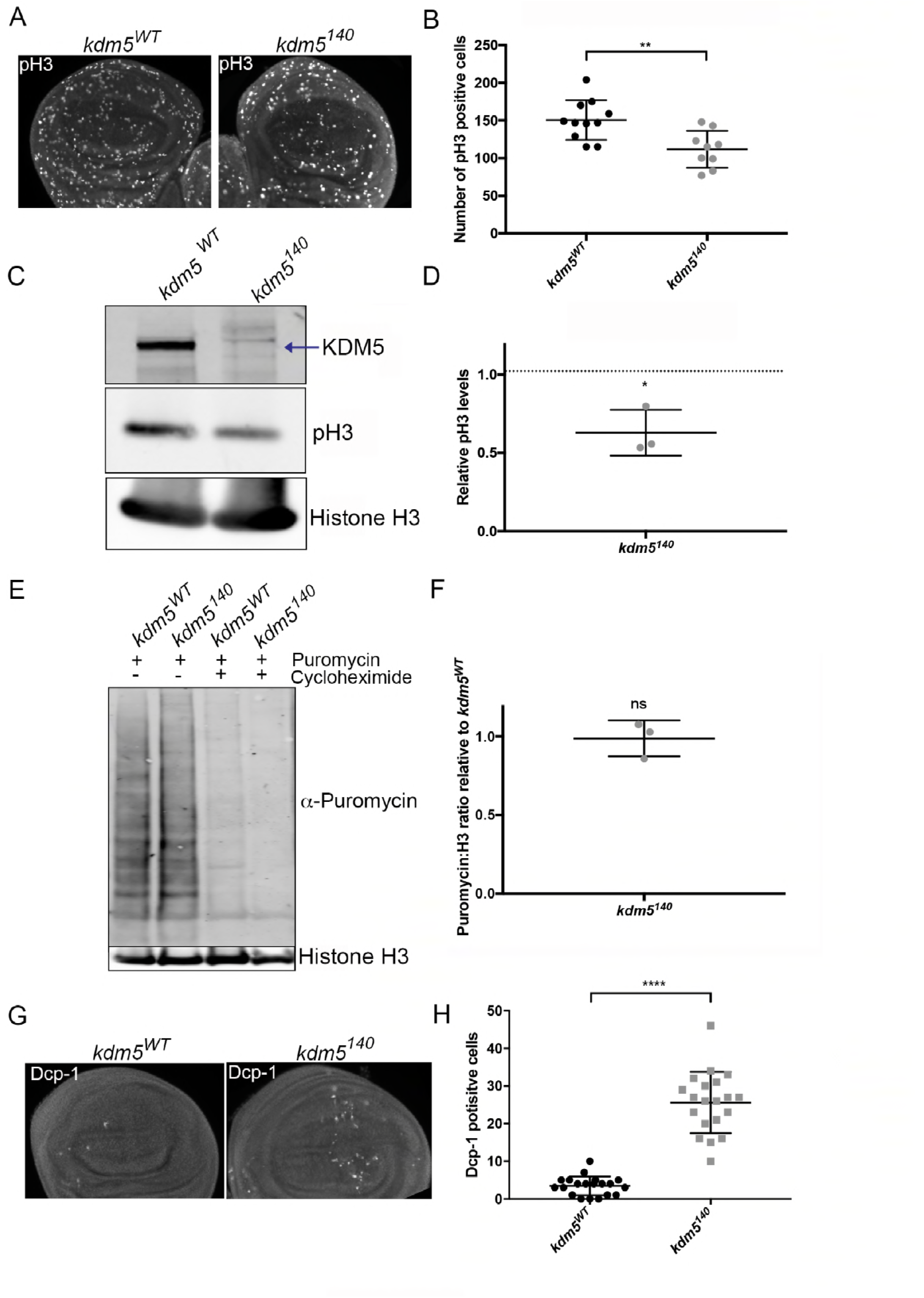
*kdm5* mutant wing discs show reduced proliferation and increased cell death. (A) Anti-phospho-histone H3 labeling of *kdm5*^*WT*^ (left) and *kdm5*^*140*^ (right) mutant wing discs matched for developmental stage (3^rd^ instar). (B) Quantification of the number of phospho-histone H3 (pH3) cells in the wing pouch of *kdm5*^*WT*^ (N=11) and *kdm5*^*140*^ mutant (N=9) wing discs. ** *p*=0.003. (C) Western blot of *kdm5*^*WT*^ and *kdm5*^*140*^ mutant wing discs (ten discs per lane) showing anti-KDM5 (top; arrow indicates KDM5 band), phospho-histone H3 (pH3; middle) and histone H3 load control (bottom). (D) Quantification of three Western blots showing increased phosphorylated histone H3 in *kdm5*^*140*^. * *p*=0.01. (E) Anti-Puromycin Western after puromycin incorporation into *kdm5*^*WT*^ or *kdm5*^*140*^ wing imaginal discs (developmental age matched). Incorporation of puromycin is blocked by co-incubation with cycloheximide. Histone H3 levels serve as a loading control. (F) Quantification of three anti-Puromycin Western blots shown as a ratio with anti-histone H3 loading control. ns = not significant. (G) Dcp-1 staining of *kdm5*^*WT*^ and *kdm5*^*140*^ 3^rd^ instar wing imaginal discs. (H) Quantification of the number of Dcp-1 positive cells in the pouch region of wildtype (*kdm5*^*WT*^; N=18) and *kdm5*^*140*^ (N=20) wing discs. **** *p*<0.0001

Because elevated cell death could also impede wing disc growth and lead to altered larval development, we quantified levels of apoptosis using an antibody to the effector caspase Dcp-1 (Sarkissian *et al*. 2014). *kdm5*^*140*^ mutant wing discs showed an increased number of Dcp-1 positive cells, suggesting that these discs have elevated rates of cell death (Figure 4G, H). Combined, decreased proliferation and increased cell death of imaginal disc cells may contribute to the developmental delay of *kdm5*^*140*^ larvae.

### Wing imaginal discs from *kdm5* null mutants have gene expression defects

To gain insight into the gene expression defects caused by loss of KDM5, we carried out mRNA-seq from 3^rd^ instar wing discs from *kdm5*^*WT*^ and *kdm5*^*140*^ homozygous mutant larvae. Because *kdm5*^*140*^ animals are developmentally delayed, we matched these samples for developmental age based on imaginal disc size, rather than days after egg laying. These analyses identified 1630 dysregulated genes in *kdm5*^*140*^ wing discs (5% FDR), 883 of which were downregulated and 747 that were upregulated (Figure 5A; Table S2). As with other gene expression defects caused by mutations in *kdm5* across a diverse range of organisms, the changes to mRNA levels were predominantly mild, averaging a 1.5-fold change for both up and down regulated genes (Lopez-Bigas *et al*. 2008; Lloret-Llinares *et al*. 2012; Liu and Secombe 2015; Iwase *et al*. 2016; Lussi *et al*. 2016).

**Figure 5:**
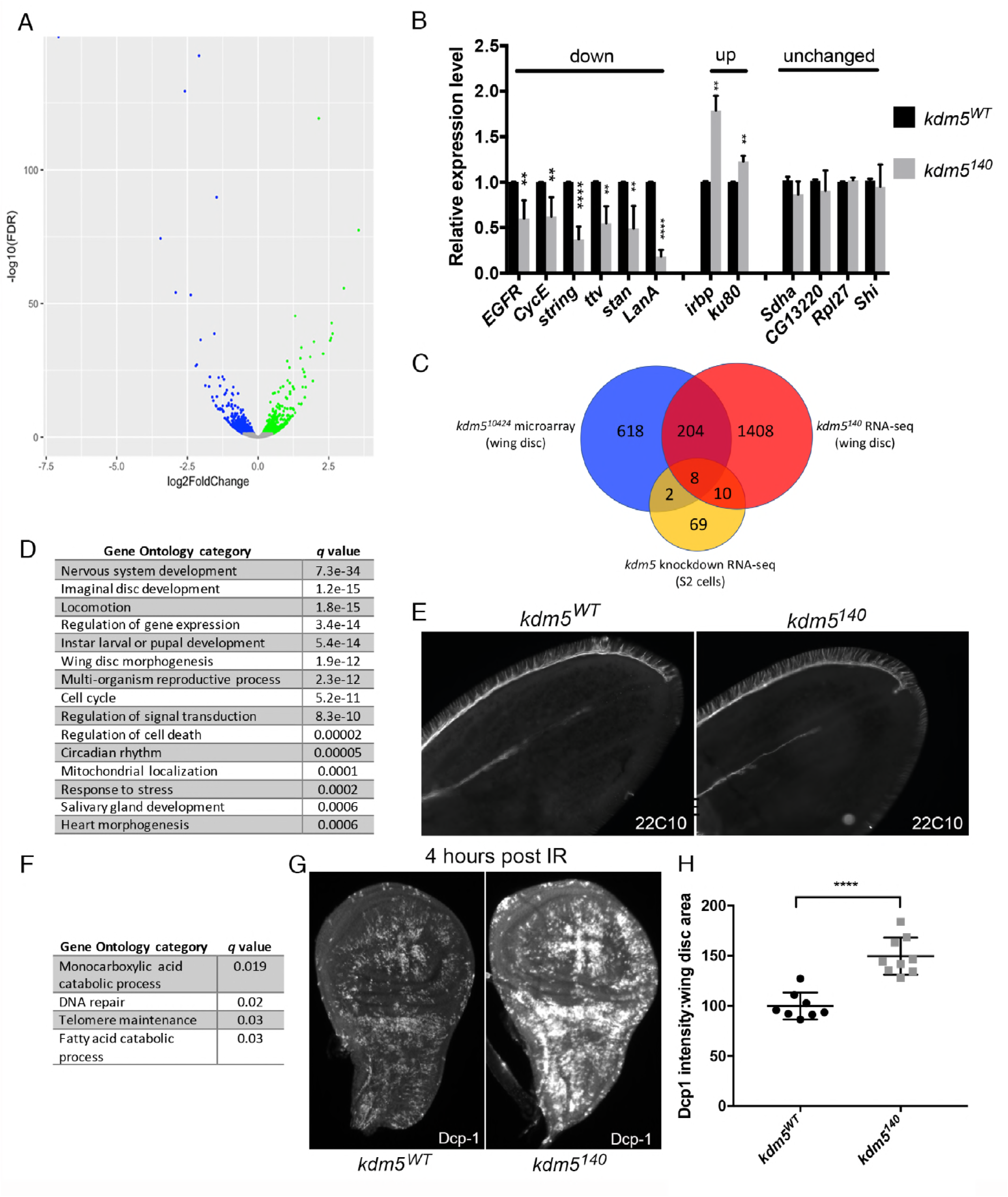
KDM5 is required for the normal transcriptional pattern of wing imaginal discs. (A) Volcano plot of genes significantly upregulated (green) and downregulated (blue) in *kdm5*^*140*^ RNA-seq data (FDR<0.05). (B) Real-time PCR validation of genes that were downregulated, upregulated or unaffected in RNA-seq data. **** *p*<0.0001, \*\*\**p*<0.001, \*\**p*<0.01. (C) Venn diagram showing overlap between current RNA-seq data and previously published microarray data from *kdm5*^*10424*^ (*p*=3.7e-48) and S2 cell KDM5 knockdown RNA-seq (*p*=0.0008). (D) Gene ontology categories enriched using Flymine (Lyne *et al*. 2007) using genes significantly down regulated in *kdm5*^*140*^. (E) *kdm5*^*WT*^ (left) *kdm5*^*140*^ (right) pupal wings stained 30 hours after puparium formation with the 22C10 antibody. (F) Gene ontology categories enriched using Flymine (Lyne *et al*. 2007) using genes significantly up regulated in *kdm5*^*140*^. (G) 3^rd^ instar larval wing imaginal disc four hours post irradiation showing cell death using anti-Dcp1 in *kdm5*^*WT*^ (left) and *kdm5*^*140*^ (right). (H) Quantitation of Dcp-1 intensity in *kdm5*^*WT*^ (N=8) and *kdm5*^*140*^ (N=9) wing imaginal discs. \*\*\*\**p*<0.0001.

To confirm the robustness of the *kdm5*^*140*^ RNA-seq, we first confirmed gene expression changes by real-time PCR of downregulated, upregulated and unaltered genes (Figure 5B). In addition, we compared our new data to our previously published microarray data generated using wing imaginal discs from *kdm5*^*10424*^ hypomorphic mutant larvae (Liu *et al*. 2014). Despite differences in platform and allele severity, 26% of genes identified as significantly dysregulated in the wing disc microarray data were similarly affected in *kdm5*^*140*^ (212/824; *p*=3.7e-48; Figure 5C; Table S3). In contrast, we do not observe significant overlap between the current RNA-seq or previous microarray data with published microarray data from wing disc *kdm5* knockdown (Figure S2) (Lloret-Llinares *et al*. 2012). It should be noted, however, that these comparisons were limited by the small number of genes that were identified as dysregulated in *kdm5* knockdown wing discs. RNA-seq data are also available from KDM5 knockdown cultured S2 cells, (Gajan *et al*. 2016), which are a macrophage-like lineage. We therefore also determined the extent to which KDM5-regulated genes in S2 cells overlapped with the changes observed in mutant wing discs *in vivo*. Perhaps unsurprisingly given the difference in cell type and context, the overlap between *kdm5*^*140*^ and S2 cell knockdown data was more modest (*p*=0.0008; Figure 5C; Table S4). Similar to previous observations, these data are consistent with KDM5 regulating distinct targets in different cell types and at different stages of development (Liu and Secombe 2015).

To define pathways affected in *kdm5*^*140*^ mutants, we carried out gene ontology (GO) analyses of genes dysregulated in *kdm5*^*140*^ using GOrilla (Eden *et al*. 2009) and FlyMine (Lyne *et al*. 2007). This revealed a large number of significantly enriched terms for downregulated genes that are summarized in Figure 5D and provided in full in Table S5 (*q*≤0.01). The reduced proliferation observed in *kdm5*^*140*^ mutant wing discs is reflected in our GO analyses, with regulators of the cell cycle being significantly enriched. These included genes such as *cyclin E* and *cdc25* that mediate cell cycle progression (Bertoli *et al*. 2013) in addition to components of key growth signaling pathways such as the epidermal growth factor receptor (EGFR) (Lusk *et al*. 2017). In addition, consistent with previous studies of *kdm5* mutants, genes involved in circadian rhythm, mitochondrial function and stress response were also enriched (Ditacchio *et al*. 2011; Liu *et al*. 2014; Gajan *et al*. 2016). Similarly, in keeping with the observation that mutations in *Kdm5* family genes are found in human patients with intellectual disability, genes involved in nervous system development were altered (Van bokhoven 2011). These encompassed a wide range of genes implicated in the development and function of the nervous system. These included transcriptional regulators such as *groucho* (*gro*) (*Agarwal et al. 2015*), actin cytoskeletal organizers such as the Abl tyrosine kinase (Kannan *et al*. 2017), and cell-cell adhesion and communication regulators such as *fascilin1*, *fascilin3*, *tout-velu* (ttv) and starry night (stan) (Elkins *et al*. 1990; Kraut *et al*. 2001; Chanana *et al*. 2009). Because the neuronal-related genes that were downregulated did not affect a single pathway or process, the importance of their dysregulation to epithelial wing disc development is not clear. We do, however, note that neuronal lineages from the wing disc include those that go on to become the sensory micro- and macrochaete of the adult thorax. As shown in Figure 2L, the specification of these cell types is not affected by KDM5, although we cannot rule out functional deficits. To further examine the connection between KDM5 and neuronal function, we examined axonal development in pupal wing discs using the antibody 22C10. At 30 hours after puparium formation (APF), *kdm5*^*WT*^ and *kdm5*^*140*^ pupal wings show similar growth of the axon along the L3 wing vein, in addition to the correct number and position of the sensory bristles along the anterior margin of the wing (Figure 5E). While we have not identified obvious neuronal defects in the wing, it is possible mild defects exist, or that the observed gene expression defects are critical in cells of the nervous system.

Fewer GO categories were identified among the genes that were upregulated in *kdm5*^*140*^ using a less stringent *q*≤0.05 cutoff (Figure 5F; Table S6). Interestingly, the inclusion of DNA repair as an enriched category suggested that *kdm5*^*140*^ mutants have increased levels of DNA damage in the absence of any exogenous mutagens. This included genes encoding Irbp/Ku70 and Ku80 that form a complex and are required for double stranded break repair and telomere maintenance (Min *et al*. 2004; Melnikova *et al*. 2005). This also indicated that loss of KDM5 may result in increased sensitivity to mutagens such as gamma irradiation. To test this, we subjected *kdm5*^*140*^ mutant larvae to gamma irradiation and found that they have increased number of Dcp-1 positive cells compared to wildtype, indicating elevated levels of cell death (Figure 5G, H). KDM5 may therefore be required to maintain genome integrity during normal development leading to *kdm5* mutants to be sensitive to DNA damaging agents. Notably, we do not find any significantly enriched GO terms by analyzing genes found to be overlapping between the current RNA-seq analyses and previously generated microarray data. This emphasizes the power of our new analyses using a *kdm5* null allele, as this enables us to better detect the relatively small changes to gene expression that are caused by loss of KDM5.

### KDM5 directly regulates many genes dysregulated in *kdm5* mutant wing discs

The 1630 genes identified as dysregulated in *kdm5*^*140*^ mutant wing discs include direct transcriptional targets of KDM5 in addition to indirect changes. Utilizing published anti-KDM5 ChIP-seq data from 3^rd^ instar wing imaginal discs, we find that 15% of dysregulated genes had an associated ChIP peak (*p*=8.8e-16; Figure 6A; Table S2)(Lloret-Llinares *et al*. 2012). Consistent with other studies showing that KDM5 proteins can activate or repress gene expression in a context-dependent manner, this included 145 (16%; *p*=8.5e-24) downregulated and 97 (13%; *p*=4.53e-12) upregulated genes. Dysregulated genes with a corresponding KDM5 ChIP signal included genes within key GO categories such as cell cycle (e.g. *cyclin E* and *stg*) neurogenesis (e.g. *groucho*), transcription (e.g. *Enhancer of polycomb*, *E(Pc*)) and circadian rhythm (e.g. *nocte*). Interestingly, these are all genes that were downregulated in *kdm5*^*140*^ mutant wing discs. Indeed, upregulated genes such as the repair proteins *ku70* and *ku80* were not identified as direct KDM5 targets (Table S2).

**Figure 6:**
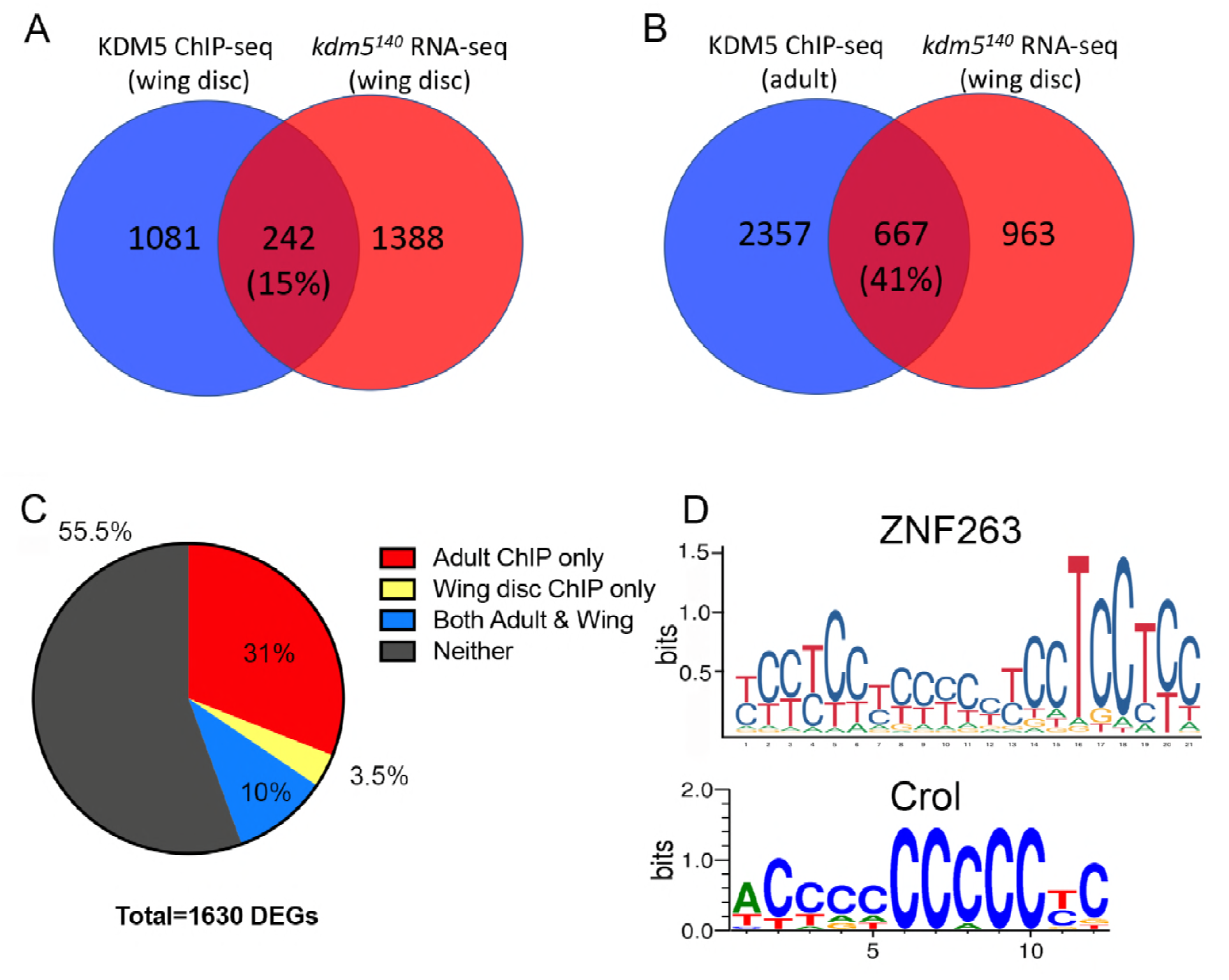
A subset of genes dysregulated in *kdm5*^*140*^ mutants are directly bound by KDM5. (A) Venn diagram showing overlap between dysregulated genes and those bound by KDM5 in ChIP-seq data from 3^rd^ instar larval imaginal discs (Lloret-Llinares *et al*. 2012). *p*=8.8e-16. (B) Venn diagram showing overlap between dysregulated genes and those bound by KDM5 in ChIP-seq data from adults (Liu and Secombe 2015). *p*=3e-144. (C) Pie chart showing proportion of 1630 dysregulated genes in *kdm5*^*140*^ wing discs and those genes bound by KDM5 in wing disc ChIP (Lloret-Llinares *et al*. 2012), adult ChIP (Liu and Secombe 2015), both ChIP datasets, and those that did not have an associated ChIP peak. See also Table S2. (D) MEME analyses of 162 directly regulated genes in wing disc and adult KDM5 ChIP-seq data showed enrichment for the zinc finger transcription factor ZNF263, which binds to the consensus sequences shown based on ChIP-seq data (Frietze *et al*. 2010). Crol is the most similar *Drosophila* gene to ZNF263 and has a similar consensus binding sequence as defined by a bacterial 1-hybrid assay (Enuameh *et al*. 2013).

The availability of ChIP data allowed us to begin addressing the key question of how KMD5 is recruited to its target genes. To investigate this, we analyzed KDM5-bound regions for the presence of the DNA sequence bound by KDM5’s ARID motif *in vitro* (CCGCCC)(Tu *et al*. 2008; Yao *et al*. 2010) in addition to binding sites for transcription factors that could mediate KDM5 recruitment. Analyses of KDM5-bound sequences using the motif analyses software MEME (Machanick and Bailey 2011) did not identify significant enrichment for the ARID binding sequence, nor for any known transcription factor binding sites (*q*<0.05). This was unsurprising since the peaks defined in the available wing disc ChIP-seq data were small (mean of 28bp)(Lloret-Llinares *et al*. 2012). We therefore also utilized KDM5 ChIP-seq data generated using whole adults to identify candidate direct targets (Liu and Secombe 2015). Despite being from a later developmental stage, 41% of genes dysregulated in the wing imaginal disc had significant KDM5 binding at their promoters using adult ChIP data (Figure 6B; *p*=3e-144; Table S2). Combining the ChIP datasets, we identified 162 high confidence direct target genes that were bound by KDM5 in the single wing disc ChIP-seq and the triplicate ChIP-seq studies carried out using adults (Figure 6C; Table S2). Interrogating these 162 KDM5-bound sequences for the known ARID DNA binding motif using MEME (Machanick and Bailey 2011) did not revealed any significant enrichment (*p*=0.6). Recognition of this sequence is therefore unlikely to be a key means by which KDM5 recognizes its target promoters. Examining KDM5-bound regions for known eukaryotic transcription factor binding factor binding sites, we identified the C_2_H_2_ zinc finger protein ZNF263 (*p*=3.1e-3)(Frietze *et al*. 2010)(Figure 6D). ZNF263 is most similar to the *Drosophila* Crooked legs (Crol) protein, which is expressed in the wing imaginal disc and has a similar *in vitro* DNA binding preference to ZNF263 (Enuameh *et al*. 2013)(Figure 6D). Interestingly, clones of *crol* mutant cells in the wing imaginal disc show reduced proliferation and increased apoptosis (Mitchell *et al*. 2008), reminiscent of the phenotypes we observe in *kdm5* mutants. Crol is therefore a candidate transcription factor that mediates KDM5 recruitment to a subset of its target genes. Testing this hypothesis awaits the identification of Crol target genes in the wing disc.

## Discussion

Here we describe the phenotypes associated with a null allele of the transcriptional regulator *kdm5*. In contrast to individual mouse knockouts of *Kdm5A*, *Kdm5B* and *Kdm5C* that are viable (Lin *et al*. 2011; Zou *et al*. 2014; Iwase *et al*. 2016), loss of the sole *kdm5* gene in *Drosophila* results in lethality. In mice, the upregulation of other KDM5 paralogs could be a key confounding factor in the analysis of individual *Kdm5* gene knockouts. For example, cells lacking KDM5B upregulate KDM5A (Zou *et al*. 2014) and KDM5B is upregulated in cells harboring mutations in KDM5C (Jensen *et al*. 2010). In a similar manner to many other genes that have more than one paralog, the phenotype(s) of true loss of KDM5 function awaits combinatorial knockout strains. Interestingly, whereas *C. elegans* also has a single *kdm5* ortholog (*rbr-2*), strong loss-of-function mutations are viable, suggesting that KDM5-regulated transcription during development may be more important in flies than in worms (Lussi *et al*. 2016). These data also point to *Drosophila* being an ideal model in which to define the essential functions of KDM5 proteins.

*Drosophila kdm5* was originally named *little imaginal discs* (*lid*) based on the size of imaginal discs in *kdm5*^*10424*^ homozygous mutant larvae (Gildea *et al*. 2000). However, our analyses of *kdm5*^*140*^ show that it would more aptly be described as a developmental delay phenotype affecting the whole animal rather than a defect only in the growth of the larval imaginal disc cells. Although they take significantly longer, *kdm5*^*140*^ mutant larvae do develop to wildtype-size before they pupate. Because pharate *kdm5*^*140*^ mutants appear morphologically normal, zygotic expression of KDM5 is not essential for the cell fate decisions required to develop external structures of the fly. Indeed, based on their appearance, it is unclear why *kdm5*^*140*^ mutants fail to eclose from their pupal case.

Interestingly, the developmental delay caused by loss of KDM5 is independent of its well-characterized JmjC-encoded H3K4me3 demethylase function. In addition, while abolishing the H3K4me2/3 binding activity of KDM5’s PHD motif mildly slows development, this chromatin recognition function does not account for the dramatically slowed larval development seen in *kdm5*^*140*^ animals. Mutant strains lacking either of these activities are adult viable, suggesting that delayed development and pupal lethality are likely to be linked (Li *et al*. 2010; Liu *et al*. 2014). These data also point to the activity of another domain or domains being critical for the developmental functions of KDM5. Previous experiments based on rescue of a *kdm5* hypomophic allele by overexpressing wildtype or domain deletion versions of KDM5 suggested that the JmjN and ARID motifs are essential for viability (Li *et al*. 2010). While these domains are good candidates for mediating key KDM5 functions, their *in vivo* activities are not clearly defined. JmjN domains are found exclusively in proteins with a JmjC motif, and the functions of these two motifs are assumed to be interdependent (Klose *et al*. 2006). This is based on crystal structure data from JmjN/JmjC domain proteins, including KDM5A, showing that the JmjN and JmjC domains make extensive contacts (Chen *et al*. 2006; Vinogradova *et al*. 2016). In addition, deletion of the JmjN motif abolishes demethylase activity in fly and mammalian KDM5 proteins (Xiang *et al*. 2007; Yamane *et al*. 2007; Li *et al*. 2010). However, while KDM5’s JmjC domain-encoded demethylase activity is not essential for development, deletion of the JmjN motif results in lethality (Li *et al*. 2010). Thus, with the caveat that deletion of the JmjN domain could have unintended structural consequences for KDM5, these data indicate that this motif could have additional, demethylase-independent, functions. Similarly, while ARID motif is assumed to have physiological DNA binding functions based on *in vitro* assays (Tu *et al*. 2008), the extent to which this occurs *in vivo* remains unclear. Indeed, structural modeling of this motif led to the suggestion that the N-terminal portion of the ARID may bind zinc to form a protein-protein interaction motif (Peng *et al*. 2015). Either the ARID or the JmjN motifs of KDM5 could, for example, interact with transcription factors such as Crol to facilitate activation of genes required for wing disc division and development (Mitchell *et al*. 2008). Much more detailed analyses of the *in vivo* functions of both of these KDM5 domains are essential to clarify their roles in development.

While nutrient deprivation can delay development (Zinke *et al*. 1999), *kdm5*^*140*^ larvae ingest food normally and do not show transcriptional changes that would indicate starvation. *kdm5*^*140*^ mutant wing discs do, however, show decreased proliferation and increased cell death. This would be consistent with previous studies showing that alleles of *kdm5* enhanced a *cyclin E* mutant eye phenotype in a manner that suggested that KDM5 is a positive cell cycle regulator during eye development (Brumby *et al*. 2004). Consistent with this occurring in more than just larval epithelial cells, cell cycle progression is also affected by knocking down *kdm5* in S2 cells that are macrophage-like cell culture line (Gajan *et al*. 2016). While the expression of cell cycle regulators was not significantly altered in S2 cells, we observed changes to numerous cell regulators in *kdm5*^*140*^ wing discs, including the G1-S phase regulator *cyclin* E and the G2-M regulator *string (cdc25*). ChIP-seq identified both *cyclin E* and *stg* as direct KDM5 targets in wing imaginal disc cells (Lloret-Llinares *et al*. 2012) but not in the adult (Liu and Secombe 2015). It is therefore possible that these cell cycle genes are bound by KDM5 only in tissues or cells that are actively growing and dividing. The regulation of cell cycle genes is also evolutionarily conserved, as KDM5A activates *cyclin E1* transcription in lung cancer (Teng *et al*. 2013).

Many factors are integrated to control the regulation of larval growth and to sense when the correct tissue and larval size has been reached for pupariation to occur. Imaginal disc size is one known determinant of developmental timing (Stieper *et al*. 2008). For example, slowing imaginal disc growth by knocking down expression of the ribosomal subunits RpS13 or RpS3 results in a developmental delay similar to that seen for *kdm5*^*140*^. Ribosomal protein genes and/or other genes required for translation were not enriched in our transcriptome analyses, nor was translation rate reduced in *kdm5*^*140*^ wing discs. Thus, while the developmental delays may be similar, KDM5 likely affects larval development through a different mechanism. One possibility for this is through the regulation of gene cycle regulators, since a hypomorphic allele of the G1-S phase regulator *cyclin E* delays larval development (Secombe *et al*. 1998). It is also possible that the proliferative changes we observe in the wing imaginal disc are an indirect consequence of a signaling defect originating elsewhere in the larva. These could include pathways such as the insulin growth regulatory pathway that do not affect developmental timing when altered in imaginal disc cells, but do when altered in the cells of the secretory prothoracic gland (Colombani *et al*. 2005; Stieper *et al*. 2008).

We also observed increased cell death in *kdm5*^*140*^ wing discs, and this occurred coincidentally with the upregulation of genes required for DNA repair. Together, these data suggest that loss of KDM5 results in higher than normal levels of DNA damage in the absence of any exogenous mutagen. Consistent with this, *kdm5*^*140*^ mutant animals were more sensitive to gamma irradiation than controls. Our previous analysis using whole larvae and a *lacZ* reporter transgene in *kdm5* hypomorphic mutant larvae also revealed an increased mutation frequency (Liu *et al*. 2014). Whether this occurs in all larval tissues, or affects some tissues more than others, remains an open question. Similar to its role in regulating key cell cycle genes, the role of KDM5 in maintaining genome stability may also be conserved in mammalian cells. Knock down of KDM5B in a range of transformed cell lines increases levels of spontaneous DNA damage in a manner consistent with defective DNA double stranded break repair pathways (Li *et al*. 2014). Thus, while the molecular mechanism remains unclear, our data provide a clear *in vivo* link between KDM5 and its promotion of cell survival by restricting DNA damage.

## Acknowledgements

The authors thank Kiera Brennan for help with initial mapping of the *kdm5* mutant in addition to members of the Secombe lab for insights at all stages of this project. Stocks obtained from the Bloomington Drosophila Stock Center (NIH P400D018537) were used in this study. We thank the Developmental Studies Hybridoma Bank (DSHB) for antibodies in addition to funding from NIH R01 GM112783, the March of Dimes (6-FY17-315), and the Einstein Cancer Center Support Grant P30 CA013330.

---

**Figure S1:**
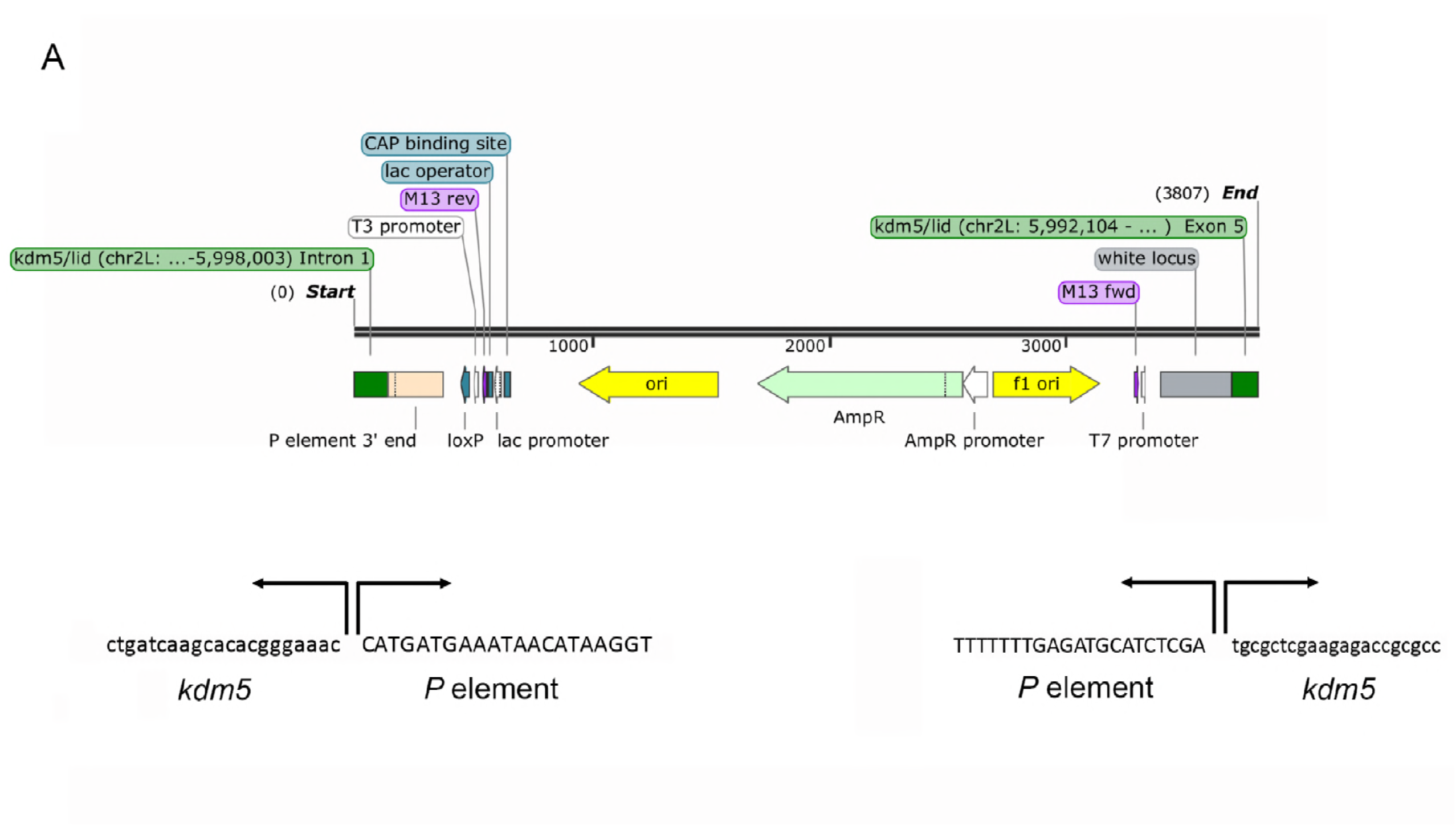
Breakpoints of the *kdm5*^*140*^ allele. (A) Imprecise excision of the NP4707 *P* element resulted in deletion that removes the 5’ end of the *kdm5* gene. The 3.8kb PCR amplicon shows the sequence surrounding breakpoints. The *kdm5*^*140*^ allele retains part of the original *P* element. Figure created using the SnapGene software.

**Figure S2:**
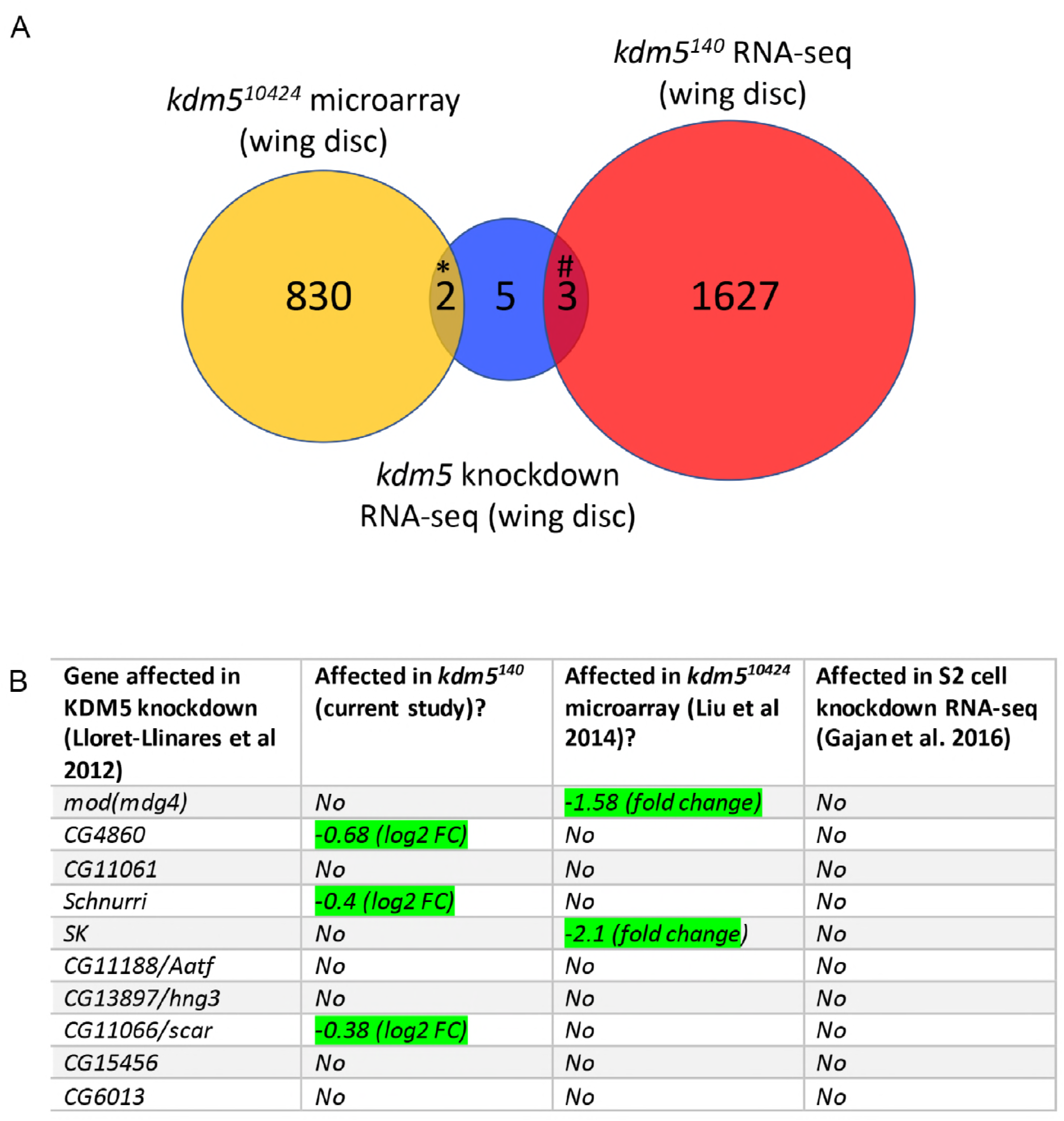
Overlap between transcriptome datasets. (A) Venn diagram showing the number of genes overlapping between current RNA-seq data using the *kdm5*^*140*^ null allele, previous microarray analyses from hypomorphic *kdm5*^*10424*^ mutants (Liu *et al*. 2014) and microarray data from *kdm5* knockdown (Lloret-Llinares *et al*. 2012). All datasets were generated using 3^rd^ instar larval wing imaginal discs. * *p*= 0.1 (not significant), # *p*=0.06 (not significant). (B) Genes significantly affected in RNA-seq from KDM5 knockdown wing discs (10 total identified using a 5% FDR), current RNA-seq data, previous microarray data and knockdown data in S2 cells. Cells labeled in green overlap between datasets.

